# Soma-to-germline BMP signal is essential for *Drosophila* spermiogenesis

**DOI:** 10.1101/2024.01.30.578090

**Authors:** Emma Kristine Beard, Rachael P. Norris, Miki Furusho, Mark Terasaki, Mayu Inaba

## Abstract

In the *Drosophila* testis, developing germ cells are encapsulated by somatic support cells throughout development. Soma-germline interactions are essential for successful spermiogenesis. However, it is still not fully understood what signaling events take place between the soma and the germline. In this study, we found that a Bone Morphogenetic Protein (BMP) ligand, Glass bottom boat (Gbb), secreted from somatic cyst cells (CCs), signals to differentiating germ cells to maintain proper spermiogenesis. Knockdown of Gbb in CCs or the type I BMP receptor Saxophone (Sax) in germ cells leads to a defect in sperm head bundling and decreased fertility. Our Transmission Electron Microscopy (TEM) analyses revealed that the mutant germ cells have aberrant morphology of mitochondria throughout the stages of spermiogenesis and exhibit a defect in nebenkern formation. Elongating spermatids show uncoupled nuclei and elongating mitochondrial derivatives, suggesting that improper mitochondrial development may cause the sperm bundling defect. Taken together we propose a new role of soma-derived BMP signaling, which is essential for spermiogenesis.

## Introduction

Bone Morphogenetic Protein (BMP) signaling is involved in a wide array of biological processes including organogenesis, immune response, and tissue homeostasis (reviewed in [1] [2]). In the *Drosophila* testis, BMP signaling from two ligands, Decapentaplegic (Dpp) and Glass bottom boat (Gbb), both secreted from the hub, is required for maintenance of germline stem cells (GSCs) [3]. These ligands signal through the type I receptor Thickveins (Tkv) and the type II receptor Punt on GSCs [3, 4]. Phosphorylation of the transcription factor Mothers against Dpp (Mad) by Tkv allows it, along with its cofactor Medea (Med), to enter the nucleus. In GSCs, Mad represses the differentiation factor Bag of marbles (Bam) to maintain stem cell identity [3].

In the testis, the role of BMP signaling outside of the stem cell niche is less understood. Recent work has demonstrated that Dpp can freely diffuse from the hub, and this diffusible fraction of Dpp signals in a different manner than contact-dependent Dpp signaling at Hub-GSC interface [5]. Diffusible Dpp signals through either of the type I receptors Tkv or Saxophone (Sax) and the type II receptor Punt on gonialblasts (GBs) and spermatogonia (SGs) to upregulate Bam expression and prevent differentiating cells from returning to the niche through de-differentiation [5]. Moreover, expression of Bam is known to regulate the number of transit amplifying SG divisions [6]. Sax, together with the downstream effector Smad on X [7] (Smox), is also found in somatic cyst cells (CCs) where it contributes to restricting over proliferation of SGs [7]. Mutant testes lacking either Sax or Smox in CCs have excessive spermatogonial division [7]. These studies have suggested that BMP signaling is not exclusively occurring between the niche and stem cells in the testis.

Soma-germline interactions in the *Drosophila* testis are vital throughout spermatogenesis, but while the pathways present in the niche and mature spermatid cysts have been well characterized, less is known about signaling between CCs and transit amplifying germ cells [8]. Direct contact between SGs and CCs is required for germline proliferation since mutations in *zero population growth* (*zpg*) and *discs large* (*dlg*), involved in gap junctions and septate junctions respectively, exhibit SG death [9, 10]. However, it is unclear what regulatory signals are exchanged at these junctions. SGs are also known to signal to CCs through the epidermal growth factor receptor (EGFR) since loss of the EGF ligand Spitz (Spi), the protease which actives Spi, Stet, and the EGFR itself lead to buildup of SGs and failure to transition to the SC stage [8, 11, 12]. Though signaling from SGs to CCs has been described, there are currently no known signaling pathways from CCs to SGs or SCs [8].

After asymmetric division of GSCs, the differentiating daughter GB undergoes four rounds of transit amplifying divisions as SGs. 16-cell SGs then become SCs and enter two rounds of meiotic divisions [13]. Throughout these stages, the germline cysts are encased by two CCs [11]. Later, after the spermatids are polarized, CCs give rise to head or tail cyst cells based on their position relative to the germ cells [11]. It is well known that soma-germline interactions are essential for proper germline development [11, 14, 15]. However, the signaling pathways occurring between the soma and germline remain largely elusive [11].

## Results

### Sax is required in the germline for proper spermiogenesis

After meiosis, syncytial cysts of 64 haploid spermatids undergo drastic changes in their nuclei (sperm heads), mitochondria, and flagellar axonemes, followed by plasma membrane remodeling and individualization [16]. Nuclear remodeling occurs through chromatin condensation facilitated by the exchange of histones for transition proteins and protamines [17]. Stages of spermiogenesis are easily identified and named by their level of chromatin condensation; round, leaf, canoe, and needle [16]. In the late canoe stage, spermatids transition to protamine-based chromatin, allowing the nucleus to form a compact needle shape [17, 18].Throughout the process of spermiogenesis, sperm nuclei are bundled together until they are transferred to the seminal vesicle [16].

Knockdown of Sax in differentiating germline cells under the bamGal4 driver caused a defect in sperm head bundling demonstrated by complete scattering of sperm heads throughout the distal half of the testis (Figure 1A, B). In the control testes, nuclei of post-meiotic spermatids were detectable as clusters from the beginning of the early elongation stage and seen as multiple bundles at the distal end of the testis (Figure 1A, C). In Sax RNAi testes, post-meiotic nuclei were still relatively clustered in the round stage of spermatids (Figure 1D). However, they were found completely scattered in leaf∼canoe stages (Figure 1B, D). Although the spermatids were able to continue maturing while unbundled, they were not able to fully complete spermiogenesis, appearing to halt at the canoe stage, and no needle stage nuclei were detected (Figure 1 C, D). Consistent with reports that protamine incorporation occurs in the late canoe stage, we found that approximately 51% (n=255 spermatid bundles) of bundled canoe spermatids in ProtamineB-GFP flies were GFP positive, while 7% (n=758 spermatids) of scattered canoe spermatids in bamGal4>Sax RNAi;ProtamineB-GFP flies were GFP positive (Figure 1F, G), indicating that Sax RNAi spermatids likely stop development during the canoe stage. Due to halted spermiogenesis, seminal vesicles were completely empty in ∼94.6% of Sax RNAi testes (Figure S1A-B, n=113) with the seminal vesicles of the remaining testes containing only a small number of sperm cells (Figure S1B’). Consistently, Sax RNAi flies showed significantly reduced fertility (Figure S1C).

**Figure 1.**
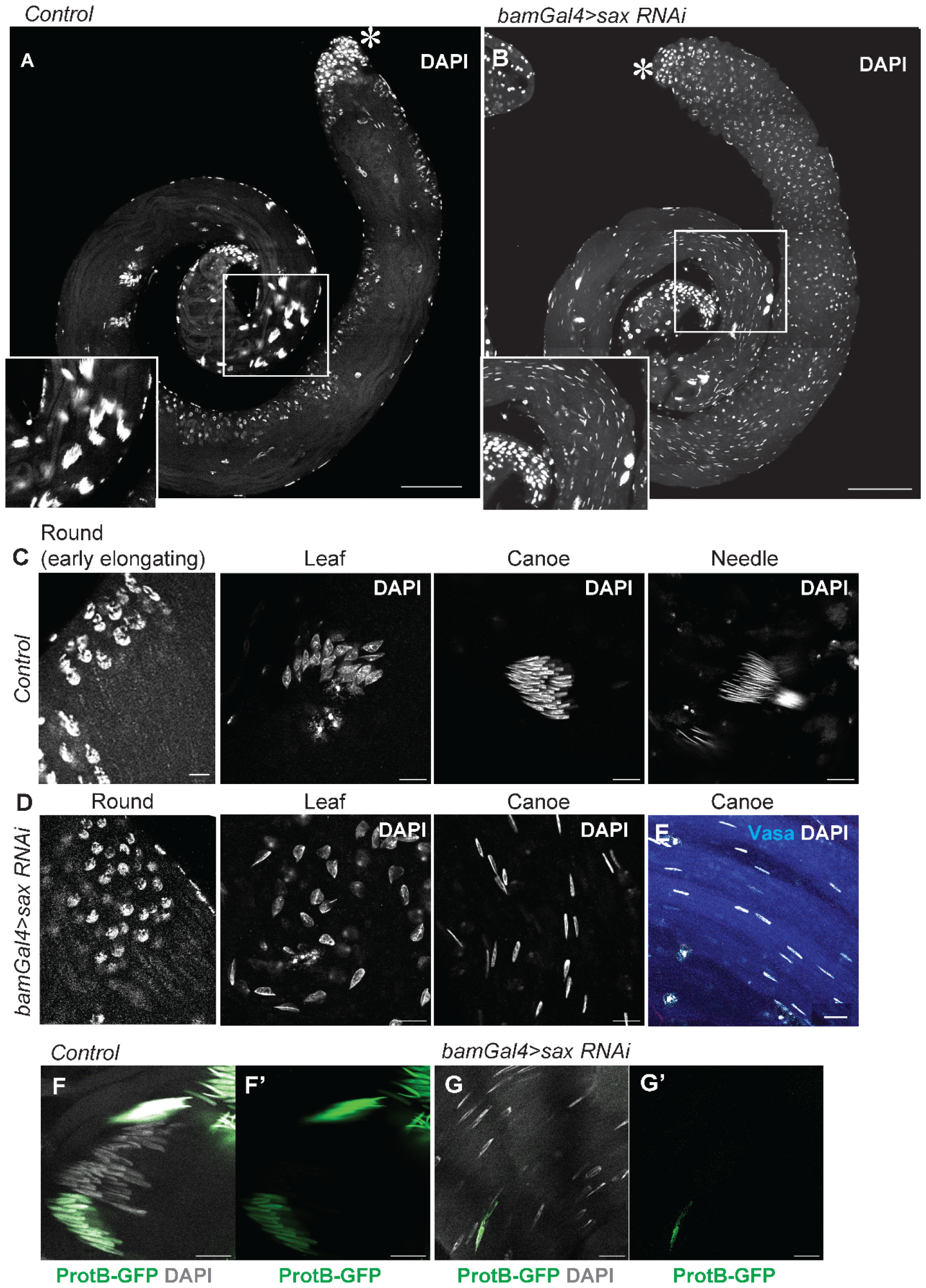
Sax is required in the germline for proper spermiogenesis. **A, B**) Representative images of testis tip without (**A**) or with (**B**) expression of shRNA against Sax under the bamGal4 driver (**B**). **C, D**) Representative images of post-meiotic nuclei of spermatids in indicated stages without (**C**) or with (**D**) Sax RNAi under the bamGal4 driver. **F, G**) Representative images of ProtamineB-GFP incorporation in spermatids/sperm nuclei in the testis without (**F**) or with (**G**) Sax RNAi under the bamGal4 driver. Scale bars in A, B are 100μm. Other scale bars represent 10 μm. Asterisks indicate approximate location of the hub.

### Soma (CC)-derived Gbb signals to the germline to maintain proper spermiogenesis

To determine which BMP ligand(s) signal to Sax to maintain spermatid bundling, we first investigated Gbb as Sax preferentially binds Gbb, and Gbb is known to be expressed in CCs in the testis [5, 19]. We knocked down Gbb in CCs under the c587Gal4 driver combined with TubGal80^ts^. After seven days of temperature shift at 29°C, Gbb RNAi testes displayed the same sperm head scattering phenotype as seen in bamGal4>Sax RNAi flies, indicating that CC-derived Gbb is responsible for this phenotype (Figure 2A-D).

**Figure 2.**
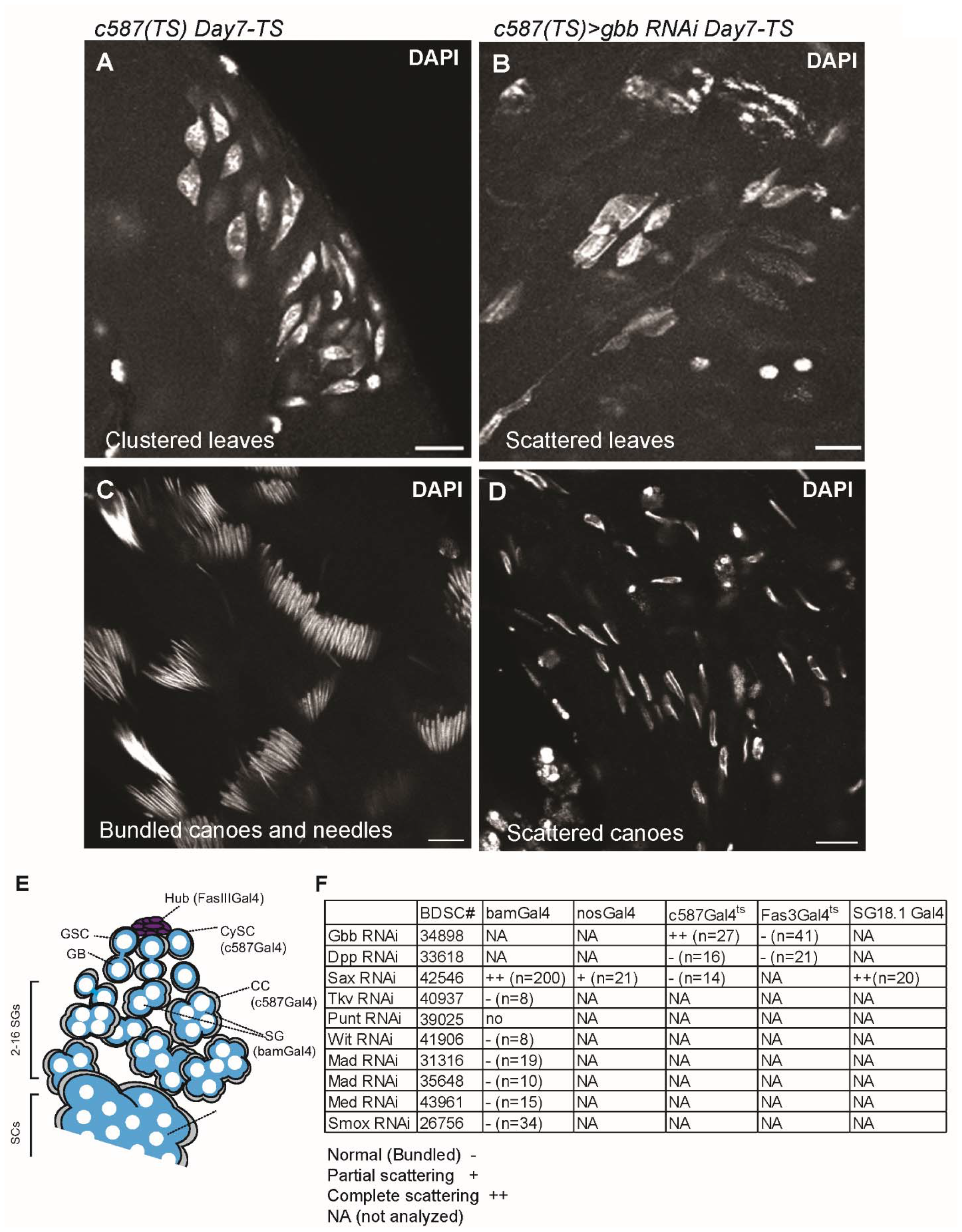
Soma (CC)-derived Gbb signals to the germline to maintain proper spermiogenesis. **A-D**) Representative images of post-meiotic nuclei of spermatids in indicated stages without (**A, C**) or with (**B, D**) Gbb RNAi under the c587Gal4 driver combined with TubGal80^ts^. Testes were dissected and analyzed for DAPI staining after 7-day temperature shift at 29 degrees. **E**) A schematic showing the cell types in the testis and expression pattern of different Gal4 drivers. **F**) Summary of sperm-head scattering phenotypes observed in indicated genotypes. Scale bars represent 10μm.

In the testes, both Dpp and Gbb, likely as a heterodimer, are required for stem cell maintenance and prevention of dedifferentiation [4, 5, 20]. However, RNAi of Dpp through either the same c587Gal4^ts^ driver, FasIIIGal4^ts^, or through DppGal4^ts^ did not lead to sperm-head scattering, indicating that Dpp is not involved in this pathway (Figure 2E, F), and that BMP signaling from CCs to developing germline cells may be solely through Gbb.

A previous study has demonstrated that BMP signal from the niche oppositely regulates Bam expression levels between GSCs and SGs, and that Sax upregulates Bam expression levels in SGs [5]. Moreover, upregulation of Bam is known to regulate dedifferentiation [5], as well as the number of SG divisions [6]. Although we did not observe severe defects in GSC number and niche appearance (Figure S2A, B), we detected excess SG division in about ∼10% of bam>Sax RNAi testes (n=22, Figure S2C, D). To further determine the cell types responsible for the observed phenotype during spermiogenesis, we analyzed additional tissue-specific knockdown of Gbb and Sax.

Importantly, when knockdown of Sax was driven by nosGal4, an early germline driver including GSCs, we observed only a subtle scattering phenotype, excluding the possibility that this phenotype is caused secondarily by a niche-GSC signaling defect (Figure 2F). We observed complete scattering of spermatids in SG18.1Gal4> Sax RNAi testes similar to bamGal4 mediated knockdown. The SG18.1Gal4 driver is expressed in late SGs and SCs, as well as the head cyst cells encasing spermatids in the final stages of maturation [21]. However, since the scattering phenotype was visible in immature spermatids, we could eliminate mature head cyst cell expression of SG18.1Gal4 in contributing to this pathway and further confirm the stage specificity of signaling. Because Sax is also reported to function in CCs [7], we examined c587Gal4 driven knockdown of Sax. These testes showed no scattering (Figure 2F), indicating that the role of Sax in differentiating germ cells to maintain spermatid bundling is not connected to its role in CCs for SG proliferation restriction.

To identify other signaling components, we screened commercial RNAi lines for the known BMP pathway core components for spermatid scattering. However, we could not successfully determine any type II receptors or known downstream effectors that caused scattering when knocked down. RNAi of the transcription factors Mad and Smox, as well as their cofactor Med, also did not show sperm head scattering (Figure 2F).

### BMP signaling controls morphology of mitochondria

Spermiogenesis involves extreme morphological changes to the spermatids to optimize the cells for fertilization. Spermiogenesis is coupled with the dynamic rearrangement of mitochondria. After the completion of meiosis, mitochondria fuse together, and form two long mitochondrial derivatives. These mitochondrial derivates wrap around each other to form the nebenkern [16].

Our Transmission Electron Microscopy (TEM) analyses showed clear morphological differences between control and Sax RNAi mitochondria. In SCs, the Sax RNAi mitochondria were larger with an average diameter at their widest point of 0.94±0.20μm (n=57) compared to 0.61±0.19μm (n=42) in the wildtype (p<0.0001, Figure 3A, B). The Sax RNAi mitochondria were also much less electron dense than wild type mitochondria (Figure 3A’, B’). Control mitochondria had tightly aligned inner and outer membranes, as well as numerous cristae (Figure 3A’). Sax RNAi mitochondria often had either a missing or broken outer mitochondrial membrane as well as very few cristae (Figure 3B’). Although the Sax RNAi mitochondria were able to aggregate and form the nebenkern, this structure was still less electron dense than in the control, and the membranes often appeared tangled together rather than concentric (Figure 3C, D). In the SG stage, some Sax RNAi mitochondria showed normal morphology and electron density, while others displayed the same low density as in SCs, suggesting that the defect begins in SGs, and that the mutant phenotype is unlikely caused by a technical artifact (Figure 3E, F). The onset of mitochondrial defects in Sax RNAi SGs is consistent with our idea that these defects are not caused by previously characterized niche (Hub)-GSC BMP signaling, but instead CC to SG signaling.

**Figure 3.**
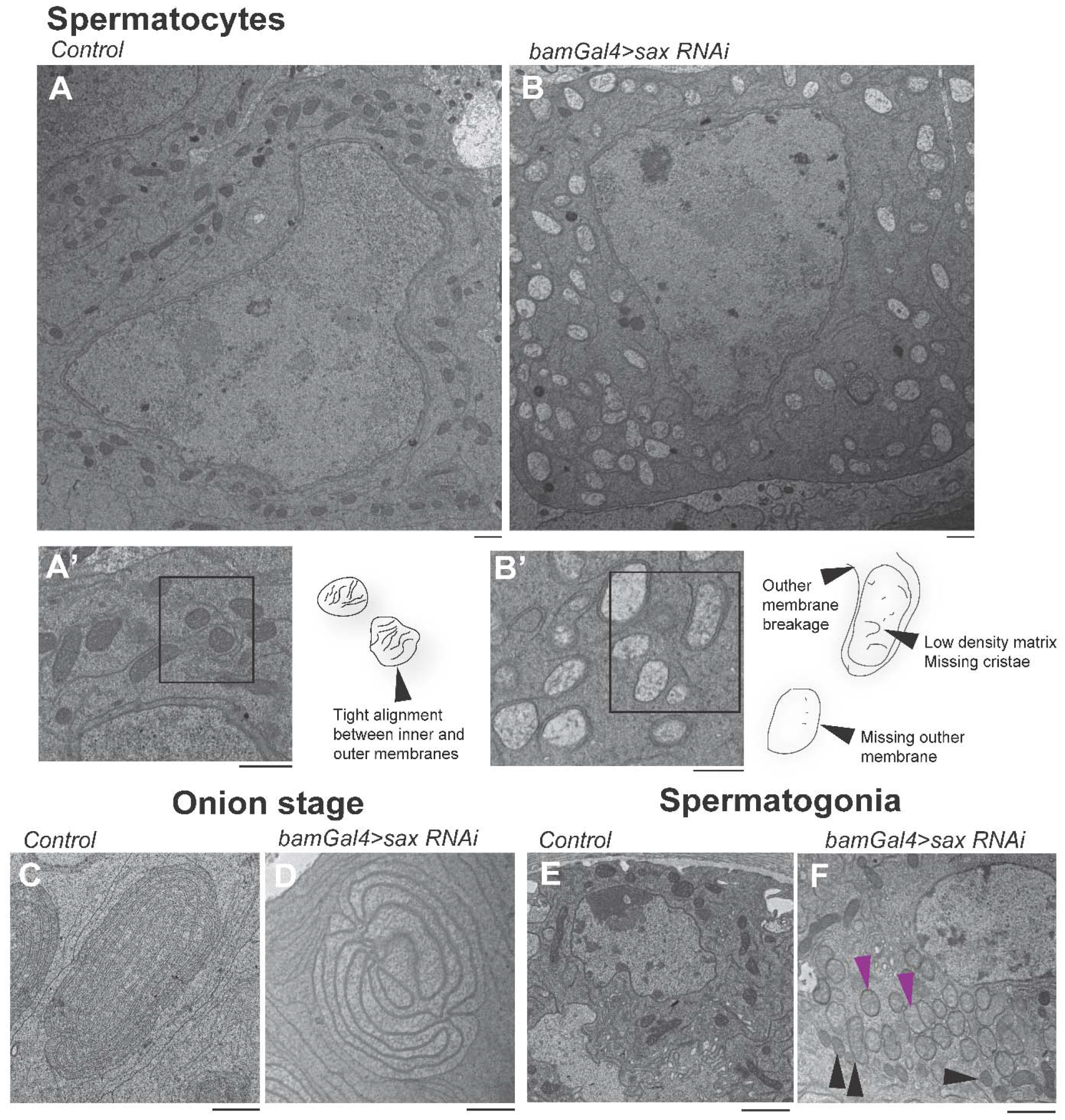
BMP signaling controls morphology of mitochondria. A) Representative electron micrographs of spermatocytes in the testes from 3- to 7-day old males of indicated genotypes. **A’** and **B’** are magnified areas in the cytoplasm showing mitochondria and graphical interpretation of their appearance. **C-F**) Representative electron micrographs of indicated cell stages in the testes from 3- to 7-day old males of indicated genotypes. Similar results were obtained by 2 more independent biological replicates. Scale bars represent 2 μm

### Mitochondria are uncoupled from axonemes and nuclei in Sax RNAi flies

In the early elongation stage, the nebenkern separates into the major and minor mitochondrial derivatives which pair with the axoneme during flagellar elongation (Figure 4A). In Sax RNAi spermatids, we observed either complete loss of mitochondrial derivatives or failure of the mitochondrial derivatives to pair with the axoneme (Figure 4B). In the late-elongation stages, individualized sperm were found in Sax RNAi, but they often contained the more than two axonemes and coupled mitochondrial derivatives were rarely found (Figure 4C, D).

**Figure 4:**
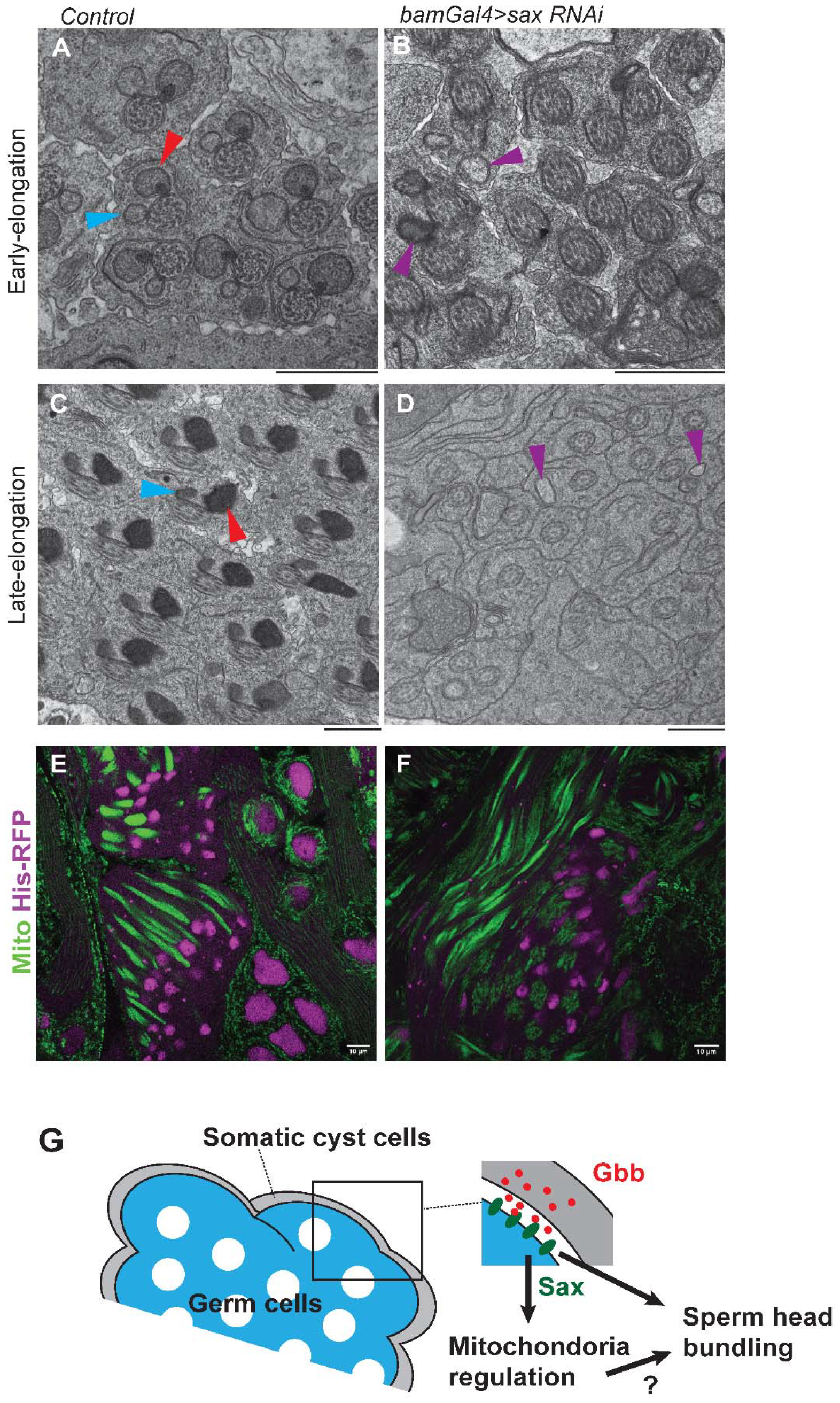
Mitochondria are uncoupled from axonemes and nuclei in Sax RNAi flies. **A-D**) Representative electron micrographs of “early-elongation” or “late-elongation” stages of spermatids in the testes from 3- to 7-day old males of indicated genotypes. Similar results were obtained by 2 more independent biological replicates. Scale bars represent 2 μm **E-F**) Representative images of live testes expressing nuclear marker, His2Av-mRFP1, together with mitochondrial marker, Tub-mito-roGFP2-Grx1, which show uncoupled nuclei and elongating mitochondria in bam>Sax RNAi spermatids. Scale bars represent 10 μm. **G**) Model. CC-derived Gbb signals to developing germline cells to regulate proper spermiogenesis. This is likely regulated through regulation of mitochondria.

What is the relationship between sperm head scattering and mitochondrial defects? In Sax RNAi testes, elongated mitochondria can be still found, but often uncoupled from nuclei and disorganized (Figure 4E, F). Previously, several sterile mutations with similar defects in mitochondrial morphology have been reported. We examined two mutant lines emmenthal (*emm*)[22] and fuzzy onions (*fzo*) [23] and found that these both show similar sperm-head scattering phenotypes (Figure S3). These results suggest a possible link between mitochondrial defects and sperm head scattering.

Taken together, we propose that soma-derived Gbb signals to Sax on SGs and SCs to ensure mitochondrial morphology and function. We suggest the possibility that proper rearrangement of the mitochondria is essential in maintaining bundling of spermatids during spermiogenesis.

## Discussion

Soma-germline interaction is known to regulate spermiogenesis in a broad range of organisms. In this study, we show that the BMP signaling pathway, a highly conserved pathway across species, is an essential component of soma-germline interaction during spermiogenesis in *Drosophila*. The ligand, Gbb, secreted from CCs, signals to differentiating germ cells through the receptor Sax. Knockdown of Gbb in CCs or the receptor Sax in germ cells leads to a defect in sperm-head bundling and decreased fertility. TEM analyses of mutant spermatocytes/spermatids revealed a morphological defect in mitochondria. Elongating spermatids showed uncoupled nuclei and elongating mitochondrial derivatives, suggesting that improper mitochondrial development may cause the sperm bundling defect. With these results, we propose the new role of BMP signaling in *Drosophila* spermiogenesis, distinct from previously characterized role of BMP signaling in the niche.

Interestingly, a previous study has suggested a possible link between BMP signal and mitochondria function. Kumar et al., identified the mitofusin2 (Mfn2) as an interactor of Smad2 and showed that TGF-ß signal regulates mitochondrial dynamics and metabolic functions through a transcription-independent role of Smad2 [25]. We found a similar phenotype in the *fzo* mutant, which is the fly homolog of Mfn2 that mediates mitochondrial fusion [23]. Although we could not determine Mad or Smox function downstream of Gbb-Sax, it would be an interesting future study to determine their function and to test direct interaction with Fzo.

## Methods

### Fly husbandry and strains

Flies were raised on standard Bloomington medium (Lab express) at 25°C (unless temperature control was required) and young flies (0- to 7-day-old adults) were used for all experiments. The following fly stocks were obtained from Bloomington stock center (BDSC); *nosGal4* (BDSC64277); *tkv RNAi* (BDSC40937); *medea RNAi:TRiP*.*GL01313* (BDSC43961); *mad RNAi:TRiP*.*JF01264* (BDSC31316); *sax RNAi:TRiP*.*HMJ02118* (BDSC42546); *punt RNAi: TRiP*.*HMS01944* (BDSC39025); *gbb RNAi:TRiP*.*HMS01243* (BDSC34898); *dpp RNAi:TRiP*.*HMS00011 (BDSC 33618); Smox RNAi: TRiP*.*JF02320 (BDSC26756). fzo[1]/TM3 (BDSC80071); emm1/CyO (BDSC11769): ProtamineB-eGFP/CyO (BDSC11769);His2Av-mRFP1 (BDSC23651). Tub-mito-roGFP2-Grx1 (BDSC67669) [26]* was used for visualizing nebenkern. *dppGal4* [27] lines are described elsewhere. *FasIIIGal4* was obtained from DGRC, Kyoto Stock Center (A04-1-1 DGRC#103-948). *bamGal4* on 3^rd^ was kind gift from Yukiko Yamashita. Temperature shift was performed by culturing flies at room temperature and shifted to 29°C upon eclosion for the 7 days before analysis. Combinations of *Tub-Gal80*^*ts*^ (a gift from Cheng-Yu-Lee) with *c587Gal4* (a gift from Yukiko M. Yamashita) were used.

### Immunofluorescence Staining

Testes were dissected in phosphate-buffered saline (PBS) and fixed in 4% formaldehyde in PBS for 30–60 minutes. Next, testes were washed in PBST (PBS + 0.2% TritonX-100, Thermo Fisher) for at least 60 minutes, followed by incubation with primary antibody in 3% bovine serum albumin (BSA) in PBST at 4°C overnight. Samples were washed for 60 minutes (three times for 20 minutes each) in PBST, incubated with secondary antibody in 3% BSA in PBST at room temperature for 2 hours and then washed for 60 minutes (three times for 20 minutes each) in PBST. Samples were then mounted using VECTASHIELD with 4’,6-diamidino-2-phenylindole (DAPI) (Vector Lab).

The primary antibodies used were as follows: rat anti-Vasa (RRID: AB_760351, 1:20; DSHB); mouse anti-Hts (1B1; RRID: AB_528070, 1:20; DSHB); AlexaFluor-conjugated secondary antibodies (Abcam) were used at a dilution of 1:400. Images were taken using Zeiss LSM800 confocal microscope with airyscan module by using 1AU-pinhole with 63X oil immersion objective (NA = 1.4). Images were processed by image J/FIJI.

Primary-secondary antibody steps were skipped for DAPI only imaging.

### Short-term live imaging

We used short term live imaging for static image acquisition to observe/quantify fluorescent-tagged proteins to avoid loss of fluorescent signal or tagged protein itself located in extracellular space by fixation and permeabilization.

Testes from newly eclosed flies were dissected into Schneider’s Drosophila medium containing 10% fetal bovine serum and glutamine–penicillin–streptomycin. These testes were placed onto Gold Seal Rite-On Micro Slides’ 2 etched rings with media, then covered with coverslips. Images were taken using a Zeiss LSM800 airyscan with a 63× oil immersion objective (NA = 1.4), with 10-20 z-stacks (interval 1μm). For all short-term live imaging experiments, imaging was performed within 30 minutes and no time-lapse imaging was performed using this method.

### Fertility assay

Individual 0- to 3-day-old males were crossed with three 0- to 3-day-old control virgin females (*yw*) in a narrow vial at 25 °C. After 3 days, males were removed. Females were left to produce embryos for 5 days. Eclosed offspring were counted for 10 consecutive days.

### Electron Microscopy

Testes were dissected into phosphate buffered saline (PBS) and then fixed in a solution of 2.5% glutaraldehyde and 3% paraformaldehyde in 0.1 M sodium cacodylate buffer on ice for 30 min. Samples were then washed in cacodylate buffer containing 2 mM calcium chloride and incubated in a solution of 1.5% potassium ferrocyanide and 2% osmium tetroxide in in cacodylate buffer, followed by washing with water and a subsequent incubation in 2% aqueous osmium tetroxide at room temperature. Samples were then washed with water and placed in 1% aqueous uranyl acetate overnight at 4°C.

The next day, samples were then dehydrated via a graded series of alcohol dilutions, then washed with propylene oxide in epoxy resin and allowed to polymerize at 60°C for 48 hours.

Ultrathin sections (60 nm) of Lowicryl HM-20-embedded testes were cut on a UC-7 ultramicrotome (Leica Biosystems) with a diamond knife (Diatome, Hatfield, PA) and imaged using Hitachi H-7650 transmission electron microscope.

### Statistical analysis and graphing

No statistical methods were used to predetermine sample size. The experiments were not randomized. The investigators were not blinded to allocation during experiments and outcome assessment. Statistical analysis and graphing were performed using GraphPad Prism 10 software. Data are means and standard deviations. The p-values (two-tailed Student’s t-test) are provided.

## Acknowledgements

We thank Margaret T. Fuller, Yukiko Yamashita, the Bloomington Drosophila Stock Center, and the Developmental Studies Hybridoma Bank for reagents; This research is supported by R35GM128678 from the National Institute for General Medical Sciences and a start-up fund from UConn Health (to M.I.).

## Author Contributions Statement

E.K.B., M.I. conceived the project, designed and executed experiments and analyzed data, drafted manuscript. M.F., R.P.N, and M.T assisted and conducted EM analyses. All authors edited the manuscript.

## References

1. Wang RN, Green J, Wang Z, Deng Y, Qiao M, Peabody M, et al. Bone Morphogenetic Protein (BMP) signaling in development and human diseases. Genes Dis. 2014;1(1):87–105. Epub 2014/11/18. doi: 10.1016/j.gendis.2014.07.005. PubMed PMID: 25401122; PubMed Central PMCID: PMCPMC4232216.

2. Sconocchia T, Sconocchia G. Regulation of the Immune System in Health and Disease by Members of the Bone Morphogenetic Protein Family. Front Immunol. 2021;12:802346. Epub 2021/12/21. doi: 10.3389/fimmu.2021.802346. PubMed PMID: 34925388; PubMed Central PMCID: PMCPMC8674571.

3. Kawase E, Wong MD, Ding BC, Xie T. Gbb/Bmp signaling is essential for maintaining germline stem cells and for repressing bam transcription in the Drosophila testis. Development. 2004;131(6):1365–75. Epub 2004/02/20. doi: 10.1242/dev.01025. PubMed PMID: 14973292.

4. Shivdasani AA, Ingham PW. Regulation of stem cell maintenance and transit amplifying cell proliferation by tgf-beta signaling in Drosophila spermatogenesis. Curr Biol. 2003;13(23):2065–72. Epub 2003/12/05. doi: 10.1016/j.cub.2003.10.063. PubMed PMID:14653996.

5. Ridwan SM, Twille A, Matsuda S, Antel M, Inaba M. Decapentaplegic ligand ensures niche space restriction inside and outside of Drosophila testicular niche. bioRxiv. 2022:2022.09.13.507868. doi: 10.1101/2022.09.13.507868.

6. Insco ML, Leon A, Tam CH, McKearin DM, Fuller MT. Accumulation of a differentiation regulator specifies transit amplifying division number in an adult stem cell lineage. Proc Natl Acad Sci U S A. 2009;106(52):22311–6. Epub 2009/12/19. doi: 10.1073/pnas.0912454106. PubMed PMID: 20018708; PubMed Central PMCID: PMCPMC2799733.

7. Li CY, Guo Z, Wang Z. TGFbeta receptor saxophone non-autonomously regulates germline proliferation in a Smox/dSmad2-dependent manner in Drosophila testis. Dev Biol. 2007;309(1):70–7. Epub 2007/07/27. doi: 10.1016/j.ydbio.2007.06.019. PubMed PMID:17651718.

8. Zoller R, Schulz C. The Drosophila cyst stem cell lineage: Partners behind the scenes? Spermatogenesis. 2012;2(3):145–57. doi: 10.4161/spmg.21380. PubMed PMID: 23087834; PubMed Central PMCID: PMCPMC3469438.

9. Tazuke SI, Schulz C, Gilboa L, Fogarty M, Mahowald AP, Guichet A, et al. A germline-specific gap junction protein required for survival of differentiating early germ cells. Development. 2002;129(10):2529–39. doi: 10.1242/dev.129.10.2529. PubMed PMID: 11973283.

10. Papagiannouli F, Mechler BM. discs large regulates somatic cyst cell survival and expansion in Drosophila testis. Cell Res. 2009;19(10):1139–49. Epub 20090623. doi: 10.1038/cr.2009.71. PubMed PMID: 19546890.

11. Schulz C, Wood CG, Jones DL, Tazuke SI, Fuller MT. Signaling from germ cells mediated by the rhomboid homolog stet organizes encapsulation by somatic support cells. Development. 2002;129(19):4523–34. doi: 10.1242/dev.129.19.4523. PubMed PMID: 12223409.

12. Sarkar A, Parikh N, Hearn SA, Fuller MT, Tazuke SI, Schulz C. Antagonistic roles of Rac and Rho in organizing the germ cell microenvironment. Curr Biol. 2007;17(14):1253–8. doi: 10.1016/j.cub.2007.06.048. PubMed PMID: 17629483.

13. Siddall NA, Hime GR. A Drosophila toolkit for defining gene function in spermatogenesis. Reproduction. 2017;153(4):R121–r32. Epub 2017/01/12. doi: 10.1530/rep-16-0347. PubMed PMID: 28073824.

14. Tran J, Brenner TJ, DiNardo S. Somatic control over the germline stem cell lineage during Drosophila spermatogenesis. Nature. 2000;407(6805):754–7. doi: 10.1038/35037613.

15. Kiger AA, Jones DL, Schulz C, Rogers MB, Fuller MT. Stem cell self-renewal specified by JAK-STAT activation in response to a support cell cue. Science. 2001;294(5551):2542–5. Epub 2001/12/26. doi: 10.1126/science.1066707. PubMed PMID: 11752574.

16. Fabian L, Brill JA. Drosophila spermiogenesis: Big things come from little packages. Spermatogenesis. 2012;2(3):197–212. doi: 10.4161/spmg.21798. PubMed PMID: 23087837; PubMed Central PMCID: PMCPMC3469442.

17. Rathke C, Baarends WM, Jayaramaiah-Raja S, Bartkuhn M, Renkawitz R, Renkawitz-Pohl R. Transition from a nucleosome-based to a protamine-based chromatin configuration during spermiogenesis in Drosophila. J Cell Sci. 2007;120(Pt 9):1689–700. doi: 10.1242/jcs.004663. PubMed PMID: 17452629.

18. Awe S, Renkawitz-Pohl R. Histone H4 acetylation is essential to proceed from a histone-to a protamine-based chromatin structure in spermatid nuclei of Drosophila melanogaster. Syst Biol Reprod Med. 2010;56(1):44–61. doi: 10.3109/19396360903490790. PubMed PMID: 20170286.

19. Haerry TE, Khalsa O, O’Connor MB, Wharton KA. Synergistic signaling by two BMP ligands through the SAX and TKV receptors controls wing growth and patterning in Drosophila. Development. 1998;125(20):3977–87. Epub 1998/09/15. doi: 10.1242/dev.125.20.3977. PubMed PMID: 9735359.

20. Bauer M, Aguilar G, Wharton KA, Matsuda S, Affolter M. Heterodimerization-dependent secretion of bone morphogenetic proteins in Drosophila. Dev Cell. 2023;58(8):645–59 e4. Epub 2023/04/14. doi: 10.1016/j.devcel.2023.03.008. PubMed PMID: 37054707; PubMed Central PMCID: PMCPMC10303954.

21. Desai BS, Shirolikar S, Ray K. F-actin-based extensions of the head cyst cell adhere to the maturing spermatids to maintain them in a tight bundle and prevent their premature release in Drosophila testis. BMC Biol. 2009;7:19. Epub 2009/05/07. doi: 10.1186/1741-7007-7-19. PubMed PMID: 19416498; PubMed Central PMCID: PMCPMC2683793.

22. Dorogova NV, Bolobolova EU, Akhmetova KA, Fedorova SA. Drosophila male-sterile mutation emmenthal specifically affects the mitochondrial morphogenesis. Protoplasma. 2013;250(2):515–20. doi: 10.1007/s00709-012-0434-2.

23. Hales KG, Fuller MT. Developmentally regulated mitochondrial fusion mediated by a conserved, novel, predicted GTPase. Cell. 1997;90(1):121–9. Epub 1997/07/11. doi: 10.1016/s0092-8674(00)80319-0. PubMed PMID: 9230308.

24. Chesarone MA, Goode BL. Actin nucleation and elongation factors: mechanisms and interplay. Curr Opin Cell Biol. 2009;21(1):28–37. Epub 2009/01/27. doi: 10.1016/j.ceb.2008.12.001. PubMed PMID: 19168341; PubMed Central PMCID: PMCPMC2671392.

25. Kumar S, Pan CC, Shah N, Wheeler SE, Hoyt KR, Hempel N, et al. Activation of Mitofusin2 by Smad2-RIN1 Complex during Mitochondrial Fusion. Mol Cell. 2016;62(4):520–31. Epub 2016/05/18. doi: 10.1016/j.molcel.2016.04.010. PubMed PMID: 27184078; PubMed Central PMCID: PMCPMC4877164.

26. Albrecht SC, Barata AG, Grosshans J, Teleman AA, Dick TP. In vivo mapping of hydrogen peroxide and oxidized glutathione reveals chemical and regional specificity of redox homeostasis. Cell Metab. 2011;14(6):819–29. Epub 2011/11/22. doi: 10.1016/j.cmet.2011.10.010. PubMed PMID: 22100409.

27. Matsuda S, Schaefer JV, Mii Y, Hori Y, Bieli D, Taira M, et al. Asymmetric requirement of Dpp/BMP morphogen dispersal in the Drosophila wing disc. Nature Communications. 2021;12(1):6435. doi: 10.1038/s41467-021-26726-6.

