## Supplementary material for "Soma-to-germline BMP signal is essential for *Drosophila* spermiogenesis": sup

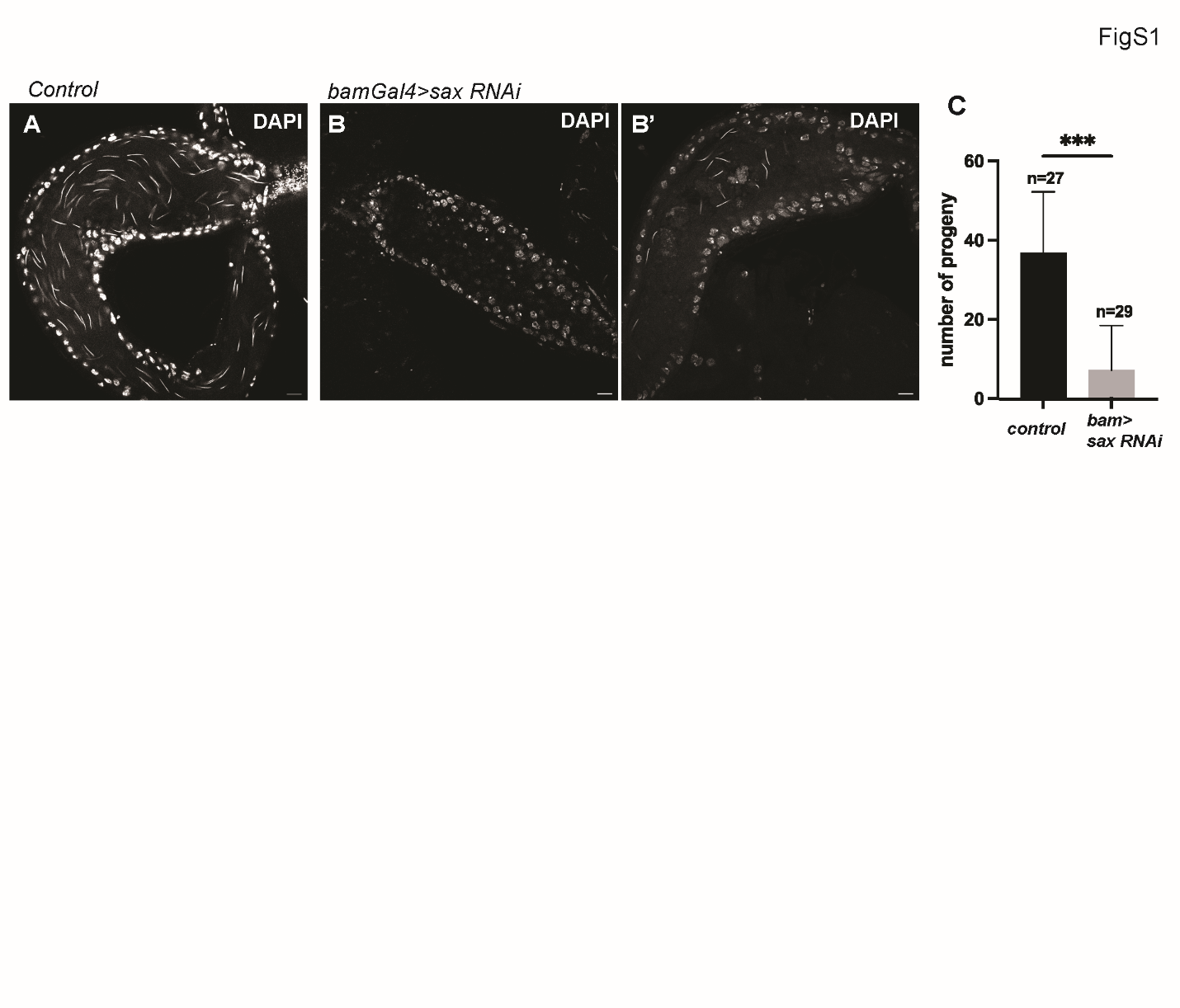

**Figure S1**

**A-B**) Representative DAPI staining confocal images of seminal vesicles of control flies (no Gal4 **A**) or bamGal4>SaxRNAi flies (**B, B’**). C) Fertility assay of Temperature shift (TS) was performed at 29 degrees for 2 days before dissection. p-values were calculated by two-sided student-t-test provided on the graph as *** (p<0.001). “n” indicates the number of tested males. Scale bars represent 10 μm.

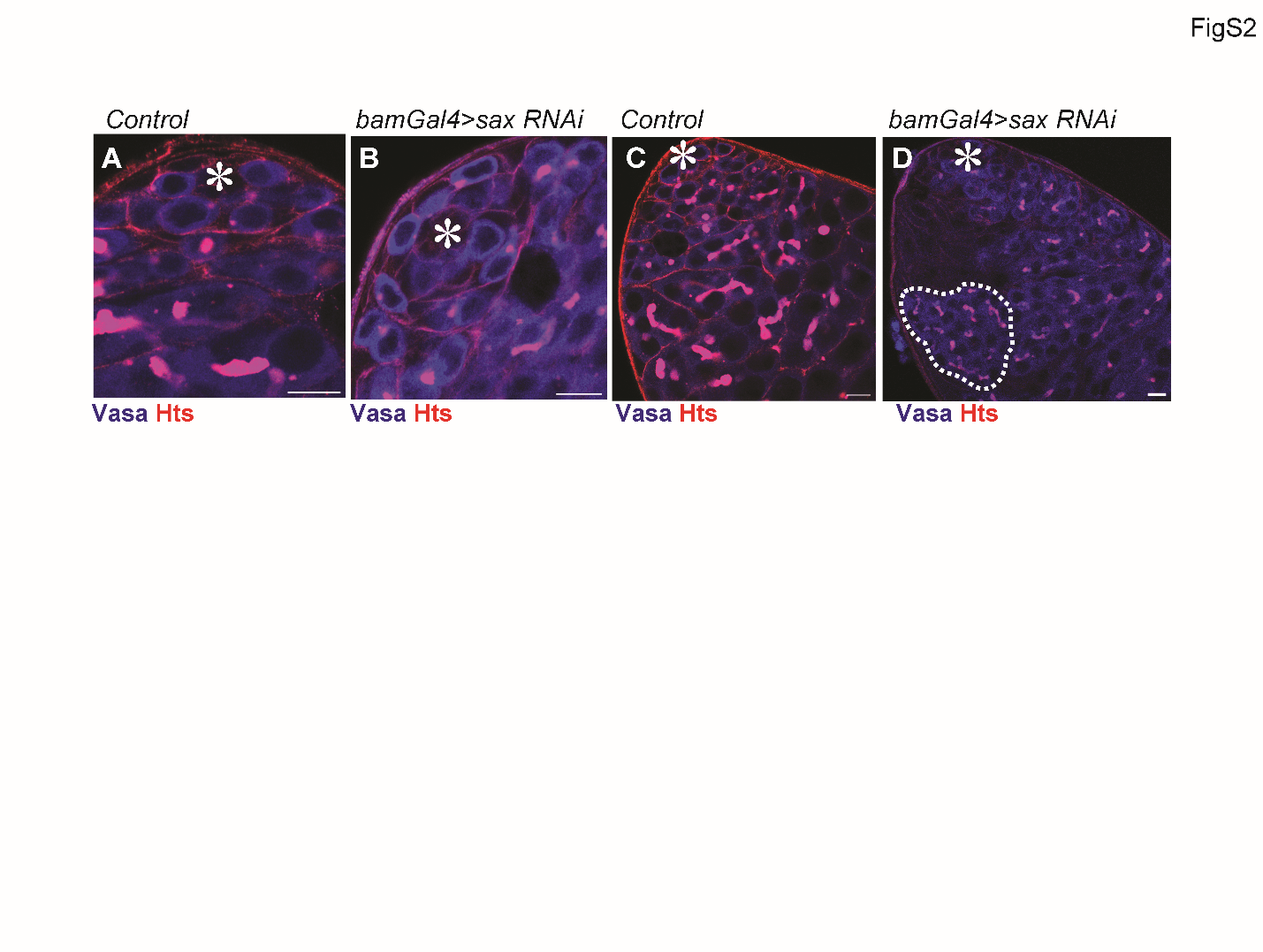

**Figure S2**

**A-D**) Representative IF images comparing testis tips isolated from control (no Gal4) fly or bamGal4>Sax RNAi flies. Scale bars represent 10 μm. The hub is encircled by red broken lines. In **D**, Sax RNAi shows >16-cell SGs indicating excess TA division (white dotted circle).

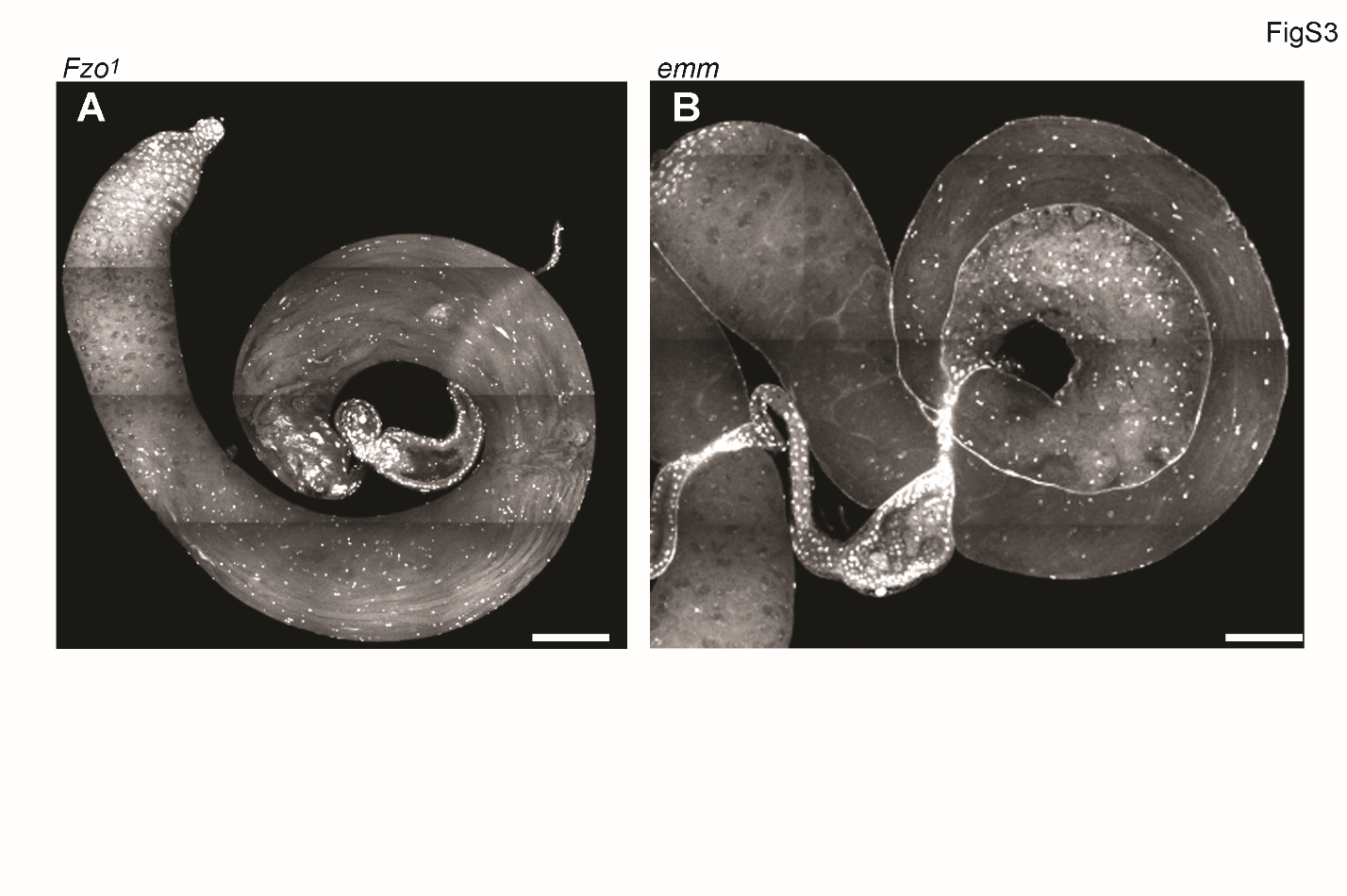

**Figure S3**

**A-B**) Representative confocal scanning images of the testes isolated from homozygous Fzo^1^/Fzo^1^ (**A**) or emm^1^/emm^1^ (**B**). DAPI (white). Scale bars represent 100 μm.
